# Repeated fentanyl abstinence intensifies opioid withdrawal and induces a proinflammatory state in striatal microglia

**DOI:** 10.1101/2025.11.06.687032

**Authors:** David J. Bergkamp, Kevin R. Coffey, Aliyah J. Dawkins, Madelyn T. Rice, Ari M. Peden-Asarch, John F. Neumaier

## Abstract

Opioid withdrawal is a serious obstacle to self-initiated abstinence, and previous experiences of opioid withdrawal may exacerbate the severity of subsequent incidences. To study the impact of repeated opioid withdrawal episodes, we compared male and female mice after one or five cycles of fentanyl exposure and withdrawal. We selectively expressed hemagglutinin-tagged ribosomes (RiboTag) in microglia of transgenic mice to immunoprecipitate and sequence RNA actively undergoing translation (the “translatome”) from striatal microglia during fentanyl withdrawal. Key changes were confirmed by RTqPCR of RiboTag RNA. Repeated bouts of fentanyl treatment and withdrawal impacted striatal microglia much more than a single cycle of fentanyl followed by withdrawal. Multiple withdrawal cycles reduced ramification of microglial processes, suggesting a more reactive cell state, and induced more severe behavioral withdrawal signs in mice. Five cycles of fentanyl exposure and withdrawal increased the expression of gene networks associated with innate immunity signaling. Indeed, 100% of the genes associated with the “microglia core sensome”, were upregulated after five cycles of withdrawal. Together these results suggest that mouse striatal microglia initiate a proinflammatory response following five, but not one, opioid exposure and withdrawal experiences and suggest that drug therapies targeting microglial innate immune responses may mitigate the severe withdrawal associated with repeated opioid tolerance and withdrawal.

**Significance statement:** Repeated cycles of fentanyl administration and withdrawal caused worsened behavioral signs of withdrawal in mice. This is the first such study to directly examine the effects of repeated opioid withdrawal on mouse behavior. We also found that repeated opioid withdrawal increased the expression of RNAs related to the proinflammatory “microglia core sensome”. Furthermore, microglia in the mouse striatum were present at a higher density with reduced ramification, which suggests that multiple opioid withdrawal experiences cause a significant change to microglial signaling-state.

---

**O**pioid withdrawal results from abstinence or use of an opioid antagonist after consuming opioids for a significant duration [1]. The aversive symptoms of opioid withdrawal vary depending on the potency, duration of administration, and clearance of each drug, with short acting opioids such as fentanyl producing very strong symptoms in the first 12-72 hours following opioid discontinuation [2]. Administration of antagonists like naloxone will precipitate withdrawal immediately leading to difficult and potentially life-threatening symptoms in certain cases [3]. Initiation of the partial agonist buprenorphine for opioid substitution therapy can also precipitate withdrawal from high potency opioids. The vast majority (86%) of opioid users seeking treatment cite opioid withdrawal sickness as their primary deterrence to initiating abstinence [4].

Opioid addiction is often framed as a “cycle” with 3 primary components: intoxication, withdrawal, and anticipation [5]. Repeated cycles are thought to induce “hyperkatifeia”, an increase in the negative emotional signs and symptoms of withdrawal. Despite this, there is surprisingly little research directly studying the effect of repeated opioid withdrawal cycles on the severity and neurobiology of subsequent withdrawal experiences. Repeated withdrawal from alcohol produces increasingly severe anxiety-like behaviors in rats [6] as well as more withdrawal-induced seizures [7] when compared to a single episode. The risk of severe alcohol withdrawal increases with the number of experiences for human patients as well. In one interesting study of human subjects with or without a history of problematic opioid use [8], those with more opioid exposure reported liking of a single opioid administration more than previously naïve individuals, however the authors saw no differences in precipitated withdrawal symptoms when each group was administered naloxone. However, they did not examine an opioid schedule that would have induced tolerance so the severity of withdrawal was not measured after chronic, recent opioid exposure. Indeed, the current preclinical literature on opioid withdrawal focuses primarily on the quantification, analysis, and treatment of a single withdrawal experience. This is incongruous with the clinical condition where chronic opioid use disorder nearly always involves numerous cycles of binge, withdrawal, and relapse. Understanding how these repeated withdrawal cycles affect the brain and subsequent withdrawal experiences is critical to developing novel therapies that may be more effective than traditional opioid replacement. Therefore, we set out to determine if repeated opioid withdrawal induces a kindling-like effect on withdrawal sign severity in mice.

A great deal of our understanding of opioid withdrawal concerns neuronal mechanisms [9] and opioid effects on specific neuronal populations [10]. However much less work has investigated how glia participate in and are affected by opioid withdrawal. There is growing evidence that mu opioid receptor expression on microglia [11] drives some of the physiological changes that occur with chronic opioid administration and contributes to opioid withdrawal [12]. Microglia also impact opioid related analgesia and tolerance by signaling via purinergic receptors [13] and toll-like receptors [14]. Previous work from our group showed that morphine withdrawal elicits major changes in microglia translation of mRNAs in the mouse striatum [15]. Therefore, we used transgenic mice that express the RiboTag gene selectively in microglia to study how repeated opioid withdrawal cycles impact the microglia translatome in striatum. In addition, we investigated striatal microglia morphology and carefully examined withdrawal behavior intensity in transgenic and wild-type mice. We found that repeated cycles of fentanyl administration and withdrawal caused worsened behavioral signs of withdrawal, increased the density and reduced the ramification of striatal microglia, and increased the expression of RNAs related to the proinflammatory “microglia core sensome”. Together these results suggest that multiple opioid withdrawal cycles cause a significant change to the withdrawal experience, and to microglial sensing and signaling-state.

## Results

### Five cycles of opioid withdrawal exacerbate behavioral signs of withdrawal in mice

We first determined how the experience of multiple cycles of fentanyl exposure and withdrawal impacted behavior. There were four experimental groups based on drug treatment: saline (Sal) or fentanyl (Fent) and number of withdrawal cycles (One or Five). Animals in the one cycle condition were injected twice daily for five consecutive days with saline or escalating doses of fentanyl (ip) and then behavioral measures were made at 16, 48 or 196 hrs following the last injection. Animals in the five cycle conditions were injected twice daily for five days and then allowed to undergo withdrawal in their home cage for four days before undergoing four additional cycles of fentanyl injection and withdrawal (Fig 1A).

**Figure 1.**
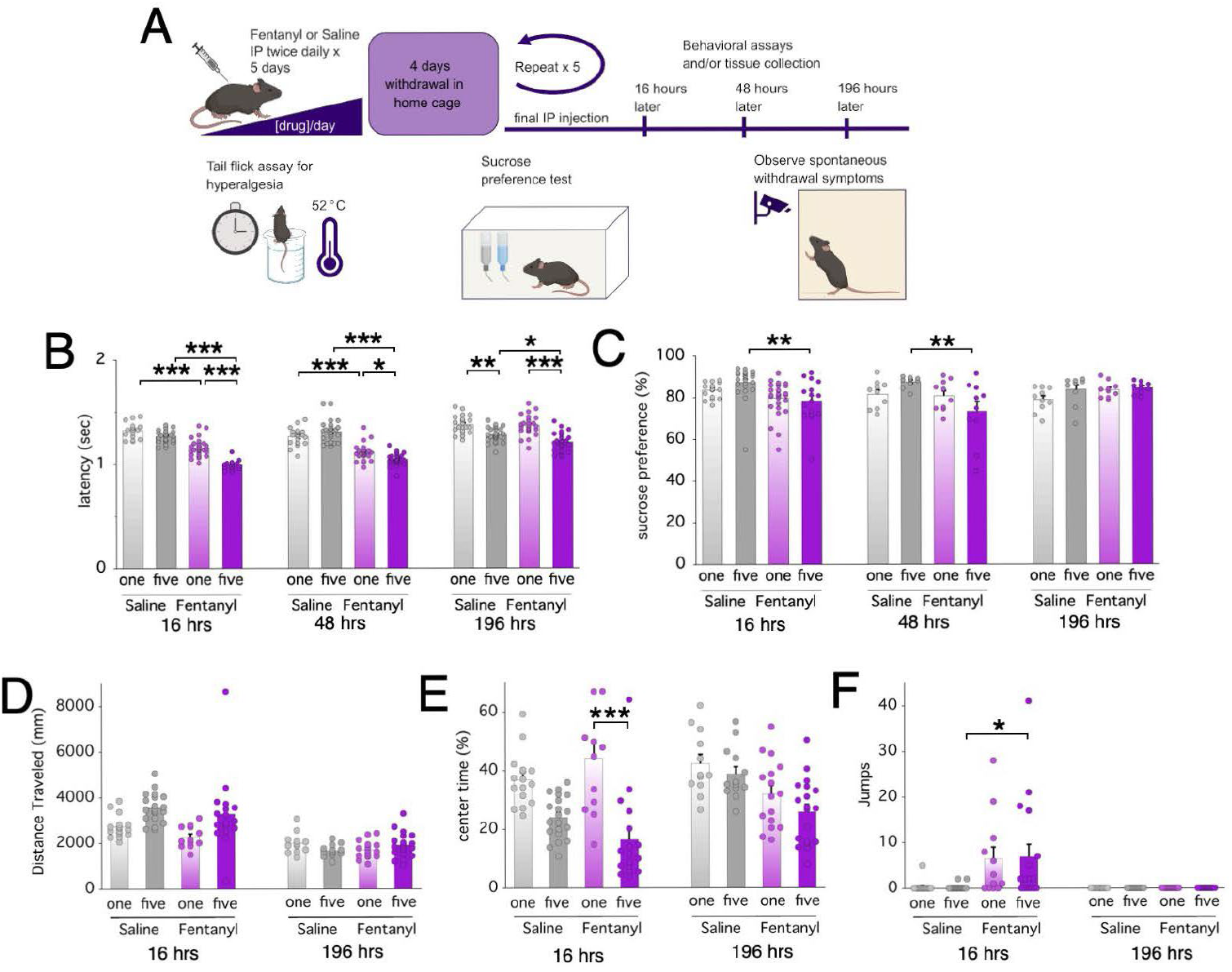
Fentanyl withdrawal symptoms are exacerbated by repeated opioid withdrawal experiences. **A**, Mice were treated with fentanyl or saline for five days followed by measurement of withdrawal behaviors. For animals in the one cycle of withdrawal group, behaviors were measured at 16, 48, or 196 hrs following the final injection on day five. For animals in the five cycles of withdrawal group, behaviors were measured at the same time points following the fifth and final fentanyl injection **B**, In the tail flick test, Fent-One (n = 28,15M 13F) and Fent-Five (n = 16, 8M, 8F) groups show significantly faster tail withdrawal latency (i.e. greater hyperalgesia) from the Sal-One (n = 16, 7M 9F) and Sal-Five (n = 23, 16M, 8F) groups at 16 and 48 hours following the last injection, but Fent-One (n = 25, 13M 12F) animals return to baseline tail flick time by 196 hours while Fent-Five (n = 27, 12M 15F) animals still show significantly faster time flick time. Note the number of Sal-One (n = 21, 10M, 11F) and Sal-Five (n = 25, 13M, 12 F) animals tested in the tail flick test at 48 and 196 hrs. **C**, Mice in the Fent-Five group had significantly reduced preference for sucrose at 16 hrs (significant effect of treatment t(79)=3.58, p=.0006) and at 48 hrs (significant effect of treatment t(36)=2.61, p=.0132 and treatment*cycle interaction t(36)=2.39, p=.0223) after the final fentanyl injection. In the Open Field Test, there was no difference between distance traveled (**D**) between groups, but Fent-Five had reduced time exploring the center of the open field (**E**), consistent with greater anxiety-like behavior. **F**, Fentanyl induced escape jumps at a significantly higher rate at 16 hrs but not at 196 hrs of withdrawal. Asterisks denote Tukey’s HSD post-hoc tests, *p<.05, **p<0.01, ***p<0.001.

We used the tail flick test to determine if hyperalgesia, normally induced during opioid withdrawal [19], was different between animals experiencing a single vs. multiple cycles of withdrawal at 16, 48, and 196 hrs after the last fentanyl administration. Fentanyl-treated mice showed hyperalgesia relative to saline treated mice, and mice with more fentanyl withdrawal experiences showed the most hyperalgesia (Figure 1B). There was no effect of sex at any timepoint with sex, treatment, and cycle as fixed effects, so the groups were collapsed for both sexes. At 16 hrs, there was a main effect of treatment (t(80)=13.97, p<.0001), cycle(t(80)=7.02, p<.0001), and treatment*cycle interaction (t(80)=3.51, p=.0007). The hyperalgesia was more pronounced for animals in the Fent-Five group compared with the Fent-One group at 16 hours following the final fentanyl injection (p<.0001) and both the Fent-Five and Fent-One groups were significantly more hyperalgesic than animals in their respective saline control groups at the 16-hour timepoint (p<.0001). By 48 hours after the final fentanyl injection, the Fent-One and Fent-Five groups continued to be hyperalgesic relative to their respective Saline controls, with a main effect of treatment (t(94)=12.27, p<.0001) and a treatment*cycle interaction (t(94)=3.45, p=.0008); the Fent-Five group was significantly more hyperalgesic than the Fent-One group (p=.02). After 196 hrs (7 days), there was a significant effect of treatment (t(95)=2.29, p=.024) and cycle (t(95)=7.41, p<.0001); the Fent-Five group was significantly more hyperalgesic than the Sal-Five (p=.0191) or Fent-One (p<.001) groups. These results demonstrate that hyperalgesia can be long lasting and more severe for animals that have experienced multiple opioid withdrawal episodes.

We used the sucrose preference test to determine if multiple withdrawal cycles induced a change in the animals’ reward sensitivity as a measure of anhedonia. We found that five withdrawal experiences led to a significant decrease in sucrose preference only for the Fent-Five group at 16 and 48 hrs after the last fentanyl injection but not for the mice exposed to a single cycle of fentanyl treatment and withdrawal (Figure 1C). The animals in the Fent-Five group drank significantly less sucrose solution than animals in the Sal-Five group at 16 hours (p=.0046) as well as at 48 hours following the final fentanyl injections (p=.0061). After 196 hours (7 days) of withdrawal, sucrose preference had returned to control levels. These data suggest that the Fent-Five animals experienced greater anhedonia and lack of motivation to consume a palatable reward compared to the Fent-One animals.

We used the open field test to determine if greater experiences of opioid withdrawal had an impact on general locomotion and anxiety-like behavior. When tested in the open field chamber for anxiety-like behavior at 16 hours after the last fentanyl injection (Figure 1D), Fent-Five animals had greater locomotion than Fent-One animals (p=.0355), but there were no significant between-group differences after 7 days of withdrawal. After 16 hrs withdrawal, animals in the Fent-Five group spent significantly less time in the center of the open field than animals in the Sal-Five group (main effect of treatment (p<.0001); the Fent-One animals were not significantly different from their comparison group (Sal-One). There were no significant differences between treatment groups after 7 days of withdrawal. We also noticed that mice in the Fent-One and Fent-Five groups showed a greater number of escape jumps from the open field 16 hours after the last fentanyl injection, but not at 7 days after the last fentanyl injection (Figure 1F), with a main effect of treatment (t(61)=3.65, p=.0005) and where Fent-Five had more jumps than Sal-Five (p=.0245) and a trend for Fent-One vs. Sal-One (p=.11). This may reflect more intense withdrawal distress in the Fent-Five animals.

Together these data suggest that mice that experienced a greater number of tolerance then withdrawal episodes also showed an increase in the severity of the withdrawal signs, especially at 16 hours following their last fentanyl dose, and longer lasting signs of withdrawal.

### Striatal microglia morphology is shifted following five but not one cycle of opioid withdrawal

We performed immunohistochemistry for Iba1, confocal imaging, and 3D reconstruction (Figures 2A-C) to determine if repeated cycles of opioid withdrawal altered morphological characteristics of striatal microglia. Microglia have highly ramified processes that interact with their neighbors to surveil for changes in activity or pathological states; they may become activated in which case their processes retract and they produce a range of immune modulators and may become phagocytic [20]. There was an interaction between treatment, cycle and sex (t(22)=2.30, p=.0312) with more microglia detected in females subjected to five cycles of fentanyl compared to one cycle only (Figure 2D), suggesting microglia proliferation in the female animals. More cycles of withdrawal appear to drive the microglia in fentanyl-treated mice toward a more ameboid, less ramified state. To quantify microglial process complexity, we calculated the ramification index (the ratio of territorial volume to cell body volume) for each cell measured. There was an overall effect of fentanyl (t(20.7)=3.42, p=.0026) and a sex*cycle interaction ((t(20.9)=2.74, p=.0125), but no relevant post-hoc comparisons were significant. Therefore, sex was collapsed and there was still an overall effect of fentanyl (t(26.3)=7.84, p=.0094); there was a trend for fent-five to have lower ramification index than fent-one (p=.0734, Figure 2E). There was a main effect of fentanyl treatment (t(20.7)=3.42, p=.0026), and an interaction between cycle and sex (t(20.7)=2.74, p=.0125), although the relevant post-hoc comparisons were not significant.

**Figure 2.**
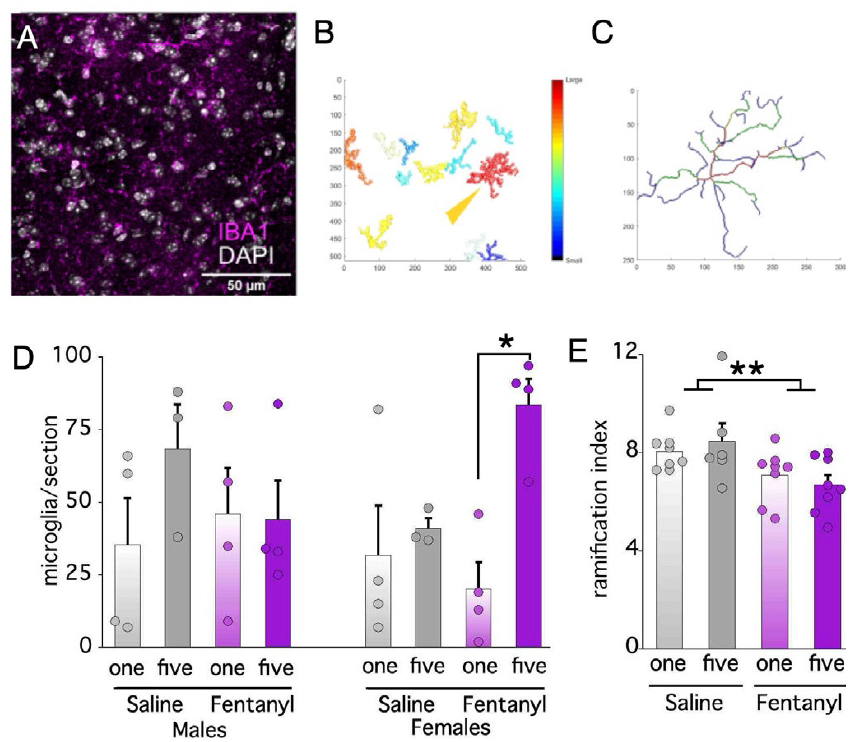
Fentanyl withdrawal reduces the ramification of striatal microglia. **A**, Maximum projection image of confocal stack showing Iba-1 staining of microglia used for morphometric analysis. **B-C**, Automated cell segmentation and removal of incomplete (border touching) cells using 3DMorph software. **D**, There were more microglia detected per tissue section in female Fent-Five compared to female Fent-One mice (* p<.05). **E**, Ramification Index was calculated as the ratio of territorial volume to cell volume for each cell. Comparing the ramification index of microglia and averaging all cells by mouse in each group (n = 8, 4 males and 4 females, in the images analyzed) via GLMM showed that there was an overall effect for fentanyl to reduce ramification (t(26.3)=7.84, ** p=.0094) but no post-hoc comparisons reached signficance.

### RNA sequencing suggests vast changes in microglia translation in mouse striatum due to repeated opioid withdrawal

Following the 16-hour time point, we dissected out the striatum (both ventral and dorsal regions) from one hemisphere of each RiboTag-expressing mouse, homogenized the tissue, and isolated the HA-tagged ribosomes to perform sequencing of the microglia-specific RNAs undergoing translation (Figure S1A). This portion, which we refer to as the immunoprecipitate (IP), represents the microglial translatome during spontaneous opioid withdrawal. We also collected a fraction of total RNA from all cells in the striatum, which we refer to as the input RNA (IN). We sequenced both IP and IN RNA from every animal to compare the microglial translatome to the whole striatum transcriptome. The sequenced RNA represents a total of 30 mice (see Table 2).

We analyzed the RNA sequencing data and determined differential gene expression using DESeq2 [21]. A principal components analysis (PCA) of all the genes showed that the IN samples separated completely from their paired IP samples along the axis of the first PC (Figure 3A). A great number of the most enriched genes in the IP samples were microglia specific (Figure 3B) and microglia transcripts were more abundant compared to other cell type genes in the IP samples (Figure 3C), which confirms that our enrichment of microglia-specific RNAs using the RiboTag method [22] was successful. Comparing IP samples by experimental treatment group suggests that the gene expression was similar for the Fent-One and Sal-One groups, which generally segregated along a combination of the first and second PC axes (Figure 3D). However, there was no clear separation between drug treatment (Fent vs Sal) groups, suggesting that a combination of withdrawal experience and drug exposure had the largest impact on the gene expression differences between groups. This was further supported when looking at the number of differentially expressed genes as determined by DESeq2. Comparing the Sal-Five to the Sal-One group, there were 45 significantly differently expressed transcripts (Figure 3E), whereas comparing the Fent-Five and Fent-One groups, we found 5503 transcripts meeting our significance thresholds (adjusted p-value < 0.01 and absolute log2(fold change) > 0.6, Figure 3F, Figure S1C). These data suggest that multiple withdrawal experiences induced more robust changes in microglia gene translation compared to a single withdrawal experience following chronic fentanyl administration.

**Figure 3.**
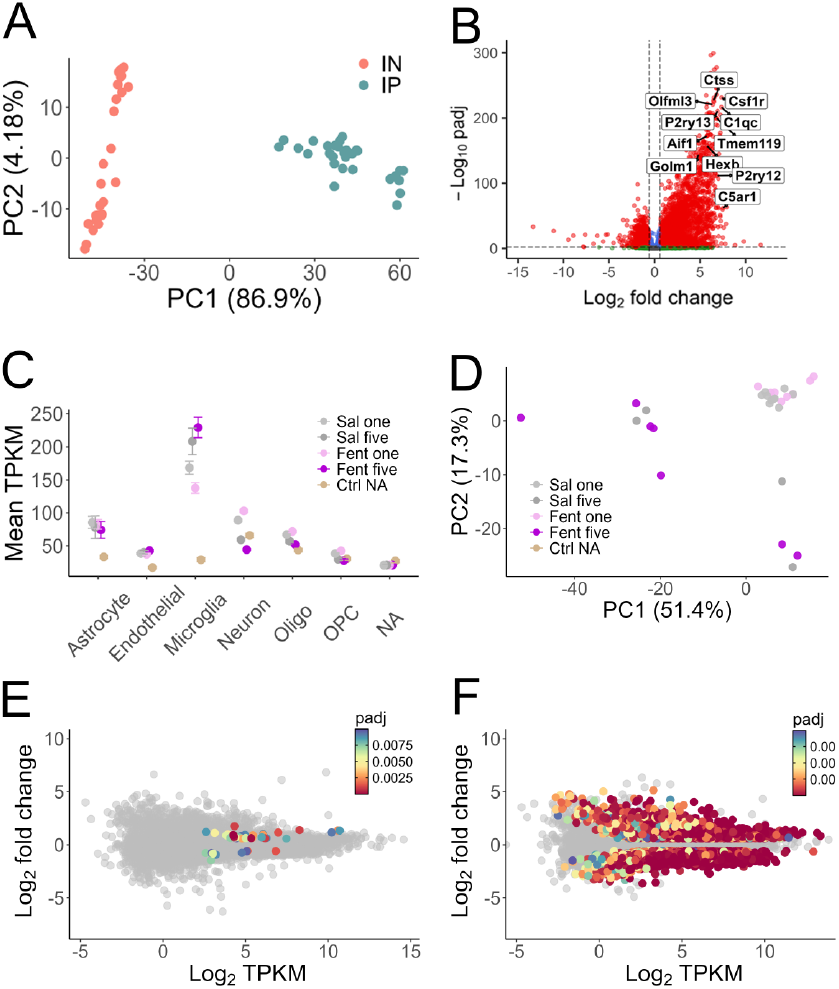
Differential expression analysis of microglia translation in the striatum shows significant effect of number of withdrawal cycles. **A**, Principal component analysis (PCA) of the input (IN) and immunoprecipitated (IP) samples’ expression together shows clear discrimination. Each point represents an individual animal. **B**, Volcano plot of adjusted p-values for each gene in IP vs IN groups versus the corresponding fold change for that gene. Comparison of IP vs IN in differential gene expression gives shows microglia genes are significantly enriched, such as Ctss, Csf1r, Olmf3, P2ry12, P2ry13, Tmem119, Aif1, Hexb, Golm1, and C5ar1. **C**, Comparison of transcripts per kilobyte of mapped reads (TPKM) in the RiboTag IP clearly indicates that microglia genes are enriched compared to transcripts typically associated with other cell types. **D**, PCA of the IP samples alone does not clearly differentiate the experimental groups, however the Sal-One and Fent-One groups do segregate from the Sal-Five and Fent-Five groups. **E**, DESeq2 analysis reveals that Sal-Five vs Sal-One yielded only 43 significantly differentially expressed genes, whereas Fent-Five vs Fent-One (**F**) found 5486 differentially expressed genes. Differentially expressed genes were defined as those with an adjusted p-value < 0.01 (method of Benjamini-Hochberg) and abs(Log2FC) > 0.6.

Following determination of differential gene expression using DESeq2, we used an unbiased approach to identify gene sets of interest in the data by applying weighted gene co-expression network analysis (WGCNA) [23]. This method determines a correlation between the expression level of each gene and every other gene sequenced. Genes are then assigned to modules based on hierarchical clustering of similar gene co-expression. These color-coded modules represent potential biological networks that can be explored further and tested with specific hypotheses. WGCNA revealed several interesting modules with overall increased mean expression of genes involved in microglia function, and when comparing the Fent-Five to the Fent-One groups, the differentially expressed genes mostly sorted into a subset of the gene modules (Figure 4A). The average Wald statistic, which accounts for the significance and direction of differential gene expression, was greater that 2.5 for five modules, indicating that many genes in these modules were robustly changed after multiple cycles of withdrawal (Figure 4B). For example, when examining the “slate” module (Figure 4C-E), the centrality of individual genes in this module was strongly correlated with increased expression in the Fent-Five vs. Fent-One groups (Figure 4D); this suggests that this module was strongly associated with the impact of multiple cycles of fentanyl tolerance and withdrawal. Overrepresentation analysis found that signaling pathways, especially GTPase-associated gene sets were significantly overrepresented in the Slate module. These proteins may play roles in signaling, cytoskeletal rearrangements, and shifting between quiescent and proinflammatory states [24,25].

**Figure 4.**
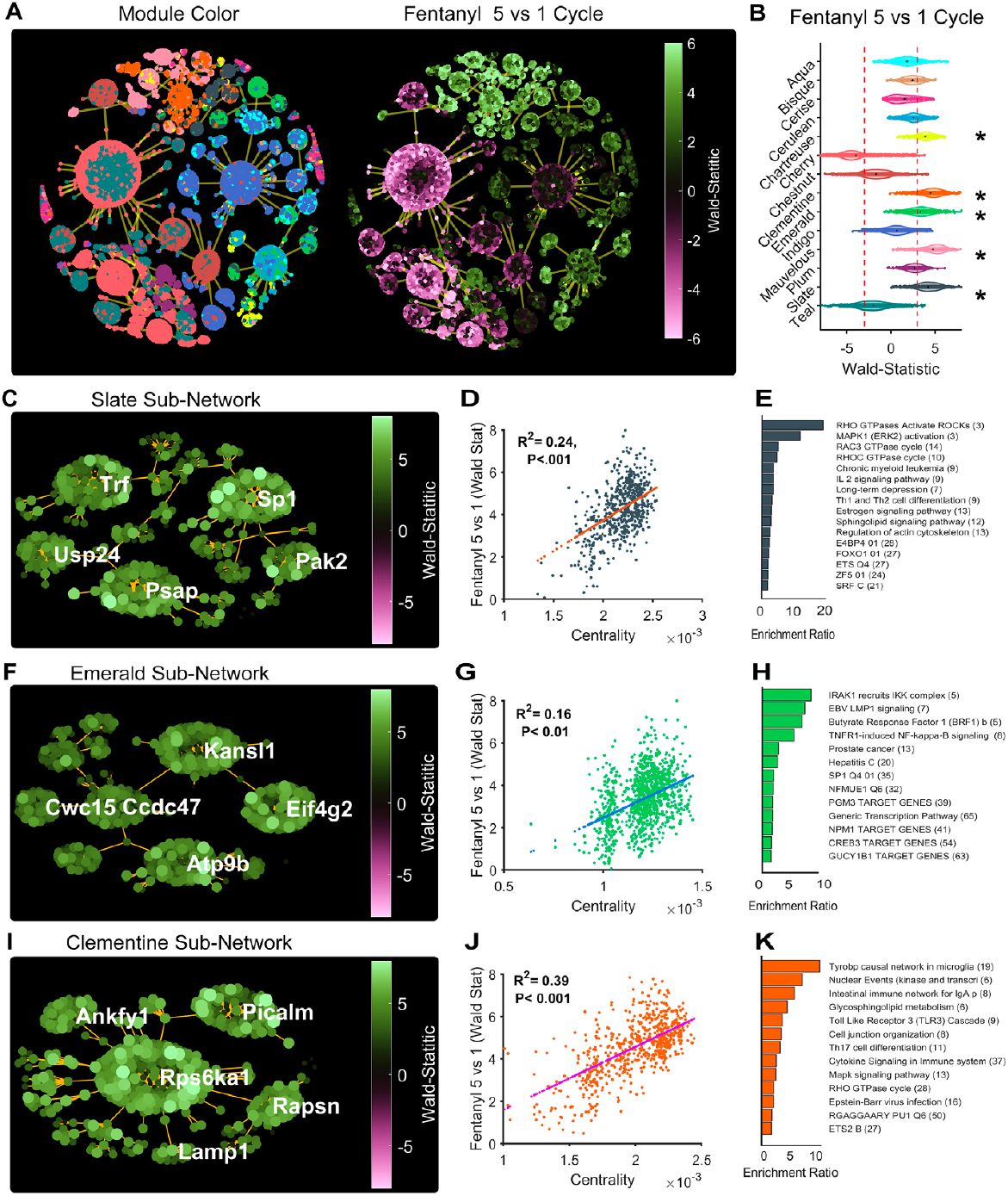
WGCNA reveals five modules of IP genes related to immune activation and microglial cell involvement in repeated opioid withdrawal: **A**, Minimum spanning trees for individual genes (as points) in the Fent-Five versus Fent-One group comparison organized into modules according to WGCNA (colored by module using R color scheme) with dashed lines indicating the genes above and below significant Wald statistic values (represents p-adjusted < 0.01 and absolute log2 Fold Change >= 2 from DESeq2 results). **B**, Violin plots of the Wald scores for all the genes within each module; five modules had a mean Wald score that was significantly increased in Fent-Five compared to Fent-One groups. **C**, Select genes in the three modules showing significant differential expression from DESeq2 results (padj from DESeq2, Benjamini-Hochberg method). Minimum spanning trees for genes in the Slate (**C**), Emerald (**F**), and Clementine (**I**) modules; the Wald statistic for each point is indicated in the colored calibration bars. **D, G, J** show the correlation of each gene’s centrality to the corresponding module and the Wald statistic for the comparison of that gene’s differential expression using DESeq2, the Pearson correlation coefficient and p values are indicated. **E, H, K**, Overrepresentation analysis on the corresponding modules was performed in WebGESTALT as described in the Methods using several databases (Keeg, Reactome, Wiki Pathways, Transcription Factor Target). The top four upregulated and downregulated gene sets from each database are displayed with the number of genes from each module that overlapped with the gene set shown in parentheses. The enrichment ratio is the number of genes from the corresponding gene set present in the module divided by the number of genes expected by chance.

The Emerald and Clementine networks contained an overrepresentation of genes associated with a range of immune and cytokine signaling processes that are associated with microglia innate immune responses (Figure 4F-K). For example, the “Tyrobp causal network in microglia” gene set had an enrichment ratio of 11.34 (p=1.35e-15). Since TYROBP (also known as DAP12) was identified as central to a microglial gene network involved in sensing microglial triggering signals (the “sensome” [26]) and includes a variety of signaling genes including purinergic receptors, cytokine receptors, and pattern recognition receptors. We then examined the 57 “core sensome” genes that are conserved in both mouse and human sensomes [27]. Surprisingly, all 57 of these genes were upregulated in Fent-Five vs Fent-One comparisons, with Wald scores ranging from 3.47-6.98. Of these, 29 were in the Clementine module, 24 were in the Mauvelous module, and 3 were in the Slate module. There was additional overrepresentation of additional upregulated genes associated with microglial activation in the Chartreuse and Mauvelous networks (Figure S2A-F).

The Cherry gene module was significantly downregulated in Fent-Five compared to Fent-One mice, and included many downregulated genes associated with neurotransmitter function (Figure S2G-I). This module also contained differentially expressed genes related to cAMP signaling and recapitulated findings of genes from our earlier study of naloxone-precipitated withdrawal from morphine [15], including seven protein kinase A regulatory and catalytic subunit genes, all of which were significantly reduced in the Fent-Five vs Fent-One comparison. It is important to note that our results here are novel and add to our previous findings, which showed increased expression of these genes when comparing morphine tolerance (animals under the influence of morphine) and acute naloxone-precipitated withdrawal.

We performed RTqPCR on a set of genes discovered by RNA sequencing to corroborate differential expression between the Fent-Five and Fent-One groups. Several genes that are strongly associated with microglia were increased in Fent-Five vs. Fent-One mice, including P2rx4, P2yr6 (in females only), CD68, and Tnfrsf1a (Figure 5A-E). There was an overall effect of cycle for Siglec1 (t(28.0)=2.24, p=.0224), but no significant post hoc comparisons (Figure 5F). Some of the RTqPCR quantifications did not replicate the DESeq2 results: e.g. Tmem119 and Pdeb3 (Figure 5G-H) did not differ significantly between groups. ß-Actin is involved in cytoskeletal reorganization that is critical to many microglial functions; it was upregulated in Fent-Five vs. Fent-One mice (Figure 5I). Together, the RTqPCR results demonstrate that most of our initial findings from the sequencing of the microglia translatome were supported by follow-up studies to validate individual gene changes.

**Figure 5.**
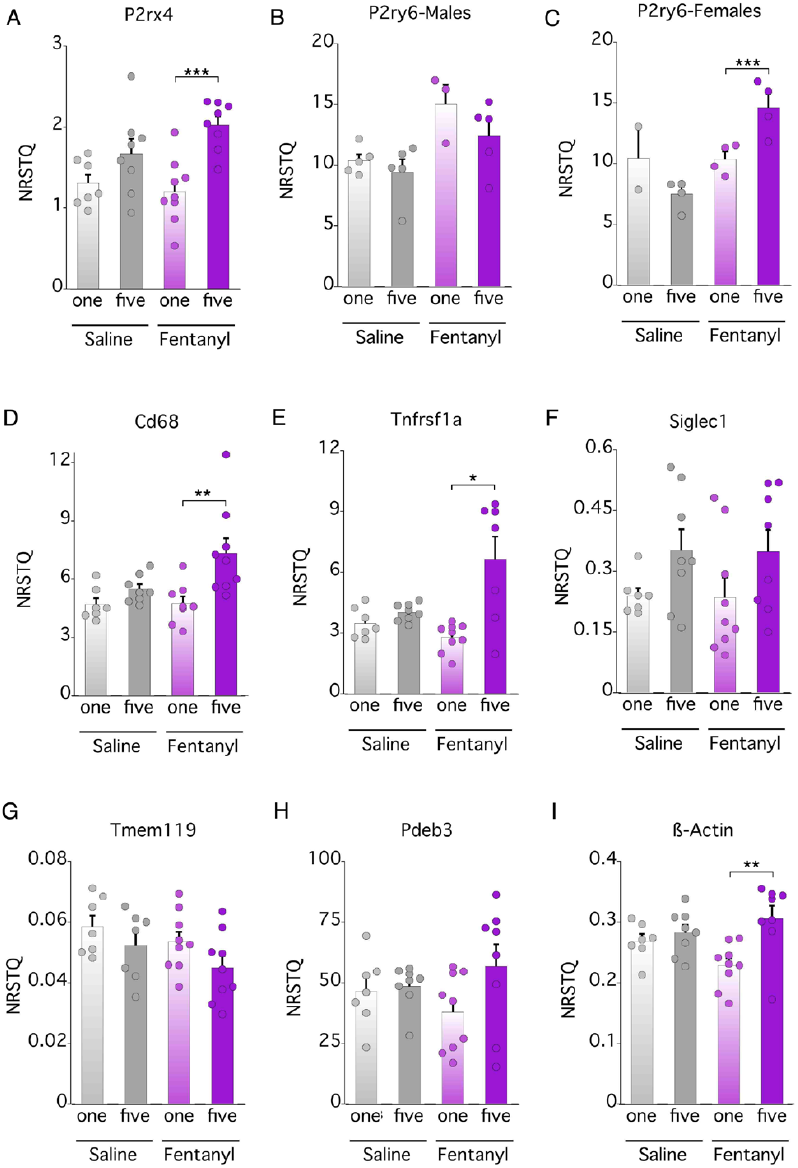
qPCR results confirmation of RNA sequencing results show that numerous genes were upregulated in Fent-Five vs. Fent-One mice. RiboTag IP RNA from individual animals was quantified by qPCR. In most cases, genes that were differentially expressed in Fent-Five vs. Fent-One in the RNAseq data were also found to be upregulated by qPCR. There was a treatment*cycle*sex interaction for P2ry6 mRNA, so the sexes were analyzed separately in this case; in all other cases there was no sex effect and the sexes were then analyzed together. **A** P2rx4, **B**), P2ry6 (males), **C**) P2ry6 (females), **D**) Cd68, **E**) Tnfrsf1a, **F**) Siglec1, **G**) Tmem119, **H**) Pdeb3, and **I**) Actb (ß-actin). Quantification was performed using standard curves as described in the Methods and data is expressed as normalized starting quantity (NRSTQ). Asterisks indicate significant effects of pairwise comparisons with Tukey’s HSD post-hoc adjusted p-values < 0.05 (*), p < 0.01 (**), and p < 0.001 (***).

## Discussion

In this study we found that five cycles of fentanyl administration and withdrawal induced a significant increase in withdrawal behaviors, indicating a greater withdrawal intensity, when compared to the same behaviors measured after a single withdrawal experience. Striatal microglia showed a reduction in ramification of their processes, which may indicate a transition into a proinflammatory state. Additionally, we found that microglia gene expression was massively altered toward a neuroinflammatory profile in the striatum of mice following five cycles of fentanyl withdrawal compared to a single cycle.

Our behavioral results show that enduring multiple cycles of fentanyl administration and withdrawal results in more severe and sustained signs of a negative emotional state. To our knowledge, this is the first set of experiments that have investigated this possibility directly. In one recent study [28], mice were administered morphine by osmotic minipump over 6 days, where one group was given saline and the other was given naloxone by injection twice daily. In this way, the experimenters induced repeated and brief precipitated withdrawal resulting in greater psychomotor sensitization to the effects of morphine and greater changes in the expression of unfolded protein response genes in the striatum, for animals given naloxone treatment. In another study [29] using rats trained to lever press for a food reward, the authors found that naloxone administration decreased responding significantly more when the rats had previously received four episodes of morphine treatment followed by naloxone. They argued that this repeated experience of withdrawal by naloxone administration may induce a conditioning effect on subsequent animal behavior. There are also anecdotal clinical reports that humans experience more severe withdrawal after multiple episodes of opioid use and abstinence. However, there is little published, empirical evidence of these symptoms and their magnitude in human literature. We propose that this worsening of opioid withdrawal symptoms over time is a general phenomenon and that it warrants further investigation in both animal models and in human subjects because it has profound implications for treating opioid use disorder.

Microglial morphology measurements after five cycles of withdrawal indicated that microglia became less ramified after five cycles of withdrawal. Decreased microglia ramification is thought to be reflective of a change in microglia signaling state [30] and suggests that there are dynamic changes evolving in microglia after repeated cycles of fentanyl and withdrawal. We argue that the changes in ramification observed after five cycles of opioid withdrawal are indicative of an increased inflammatory signaling state in striatal microglial.

The RNA sequencing data from microglia indicates that repeated opioid withdrawal induces massive changes in the expression of signaling molecules associated with inflammation compared to a single withdrawal experience. Numerous purinergic receptors were significantly upregulated (P2rx1, P2rx4, P2rx7, P2ry2, P2ry6, P2ry10b, P2ry12, P2ry13); these have previously been identified as mediators of inflammatory signaling [31], [32]. We also found increased expression of many toll-like receptor genes (Tlr1, Tlr2, Tlr3, Tlr4, Tlr5, Tlr9, Tlr12, Tlr13), which contribute to inflammatory signaling. The cross-talk between increased signaling at these receptors is difficult to predict, but P2Y and toll-like receptors can mutually contribute to proinflammatory states including the formation of inflammasomes, which are important mediators of cytokine release [33]. Importantly, these changes were not present in our Fent-One versus Sal-One nor our Sal-Five versus Sal-One comparisons, indicating that these inflammatory gene expression changes are reflective of the five repeated opioid withdrawal experiences on microglia signaling.

A major advantage of the RiboTag strategy is that it provides a deeply sequenced snapshot of the RNAs actively undergoing translation at the time of sacrifice. This information differs from transcriptomics, which includes non-translated RNAs and is impacted by the rate of RNA synthesis and degradation, and single cell RNAseq strategies that have lower sequencing depth but characterize potential phenotypically defined subsets of cells. We collected the RNA at a time of intense physical withdrawal from fentanyl, and it is possible that examining multiple timepoints will provide additional insights into the behavioral consequences of fentanyl withdrawal in microglia.

A potential limitation of this data set was that there may be a shared stress effect of repeated ip injections in the Sal-Five and Fent-Five animals, however the differences between Sal-One and Sal-Five RNA expression were quite distinct from those observed in comparison with the Fent-Five group. Indeed, the Sal-Five gene expression changes were mainly present in the input RNA (the total striatal transcriptome) rather than the IP (microglial translatome), indicating that multiple cycles of saline injection were more likely involving non-microglial cells such as neurons (FigS1B). There were no gene expression changes between groups based on animal age which is important to note as age may affect microglia gene expression [34].

We detected decreased expression of cAMP-related genes in microglia after five cycles of withdrawal (see supplementary materials for all genes in the Cherry module). In a previous investigation [15], we detected a decrease in cAMP-related genes during morphine exposure and an increase in expression 2 hrs after naloxone-precipitated withdrawal. These results suggest that the cAMP related changes in microglia occurring after a single withdrawal experience may become blunted after repeated withdrawal experiences.

The interpretation of microglial morphology and gene expression patterns to the phenotypic progression from a homeostatic to an activated state is a rapidly evolving subject, and while we monitored withdrawal-associated behaviors over an extended interval, our measurements of microglia morphology were restricted to fewer timepoints. We were struck, however, that all 57 RNAs associated with the “microglial core sensome” [27] were significantly upregulated in the Fent-Five group compared to the Fent-One group. Furthermore, we found that the “Tyrobp causal network” was significantly overrepresented in the Clementine network of the WGCNA results, and Tyrobp signaling is considered a central hub of the microglial sensome [26]. The Tyrobp gene encodes the protein TYROBP (also known as DAP12), a membrane-associated microglial protein that mediates immune signaling by coupling membrane receptors with intracellular effectors such as Src kinases [35] and is thought to play an important role in the phenotypic switch from homeostatic (quiescent) to proinflammatory states. To our knowledge, this is the first report of the involvement of these signaling processes in drug withdrawal.

Previous reports indicate that opioid exposure and withdrawal are associated with induction of inflammatory signaling. For instance, chronic morphine exposure induces microglial expression of inflammation related genes in rat spinal cord [36]. Another study [37] showed that direct opioid agonist delivery to the nucleus accumbens of rats induced local release of cytokines (IL-1α, IL-1β, and IL-6) and greater expression of microglia Iba1 protein in the ventral tegmental area. In opioid withdrawal, levels of inflammatory markers such as NF-kB can be detected even after long periods of abstinence from opioids [38]. In human subjects both alcohol use disorder [39] and opioid use disorder [40] have been linked to increases in CNS inflammation. Indeed, other authors have suggested that microglia may be central actors in promoting neuroinflammation following chronic opioid administration [41] and our work suggests this may require repeated periods of opioid withdrawal.

Our work strongly implies that microglia in the mouse striatum are driven into an inflammatory signaling state involving release of interferons after multiple cycles of fentanyl exposure and withdrawal. This process likely involves release of cytokines, and purines that activate microglial receptors included in the sensome. We found increased expression of Il6ra in our Fent-Five versus Fent-One, suggesting that IL-6 may play a role in the microglial response to repeated opioid withdrawal. Activating striatal microglia using Gq-DREADDs has been shown to induce release of IL-6, producing conditioned place aversion driven by activity at IL-6 receptors on microglia [42]. Another important component of the microglial inflammatory response is the NLRP3 inflammasome, which is mediated by TLR4 signaling and STAT1 transcription factor activity [43]. Stat1 and Tlr4 transcripts were both increased in our Fent-Five versus Fent-One sequencing data, which again supports the notion that increased microglial inflammatory signaling is a result of repeated opioid withdrawal.

In summary, the changes morphology and genes actively undergoing translation observed after five cycles of fentanyl administration and withdrawal are indicative of an increased inflammatory signaling state in striatal microglia. Combined with our behavioral results, these findings suggest that therapies that can mitigate proinflammatory signaling in microglia may be particularly impactful in treating clinical populations who regularly experience repeated cycles of fentanyl consumption and withdrawal and may complement existing strategies which are insufficient in chronically relapsing patients who are at the greatest risk for dangerous outcomes from opioid addiction.

## Methods

### Animals

Transgenic mice were generated by crossing tamoxifen-inducible Cx3cr1-CreERT2 hemizygous mice (Cx3cr1-Cre+/−, Jackson laboratories, #021160 extensively backcrossed to C57BL/6J mice) with homozygous, floxed RiboTag mice (RiboTag+/+, bred in house from a continuously backcrossed line). Male and female mice received tamoxifen (75 mg/kg dissolved in corn oil) via intraperitoneal injection (ip) for 5 days starting at 4 weeks of age to induce Cre-mediated recombination. Males and females were separately housed (2-5 mice/cage, 18-32g and 8-14 weeks old when randomized into groups) in a temperature- and humidity-controlled vivarium with a 14–10 light–dark cycle with ad libitum access to food and water. Most experiments were performed during the light phase except for the sucrose preference test described below, which was performed overnight beginning at 5 PM over several days. All experiments were performed in compliance with the Guide for the Care and Use of Laboratory Animals (NRC 2011) and were approved by the Institutional Animal Care and Use Committees for both the University of Washington and for the Veteran’s Affairs, Puget Sound Health Care System. Hemizygous (RiboTag+/−::Cx3cr1-Cre+/−) offspring were used for RNA immunoprecipitation and sequencing while Cre-null (RiboTag+/−::Cx3cr1-Cre-/-) cage mates were perfused for immunohistochemistry (IHC). Live decapitation was used for RiboTag RNA isolation procedures while cardiac perfusion was conducted for tissue preservation and performance of IHC, described below.

### Drugs

For all experiments involving fentanyl treatment (ip) injection was performed twice daily for five days to induce opioid tolerance (Fig 1A). Fentanyl citrate was supplied by the National Institute of Drug Abuse Drug Supply Program. Animals in the fentanyl treatment group were exposed to increasing fentanyl concentrations each day; the dose doubled each day, starting at 200 µg/kg on day one and increasing to 3200 µg/kg on day five. Control mice were injected with the same volume of sterile saline. Animals exposed to five cycles of withdrawal repeated the injection protocol a total of five times. Between each cycle of injections, animals in both the saline and fentanyl conditions were allowed to rest in the home cage without any injections for four days. Prior to ip injections, both fentanyl and tamoxifen solutions were sterile filtered (0.2 μm pore Whatman syringe filters).

### Sucrose preference testing

The sucrose preference test procedures were carried out following previously established methods [16]. Briefly, animals were first exposed to two 50 mL BioServ bottles (#9019) in their home cages for a period of two days. The bottles contained either water or 2% mass/volume sucrose in water. Bottles were washed daily, the contents refilled, and the side of the cage assigned to each bottle was alternated daily. After two days of training with their littermates, animals were isolated to individual cages at 5 PM the following night (3 hours prior to the beginning of the dark cycle) and again exposed to both bottles. Masses of the bottles were recorded, and mice were allowed ad lib access to the bottles until 9 AM the following day (16 hours total). The following morning, bottle masses were again recorded, and the difference was determined.

### Tail flick testing

The tail flick testing procedure was performed following previously reported methods [17]. Briefly, animals were first handled for at least three days before any measures were taken. Using a hot plate, a beaker filled to 1L with water was brought to 52°C while under constant stirring with a stir bar. Mice were then held gently in the experimenter’s hand, using a disposable paper towel as a restraint, and their tails submerged into the water bath. The time for each animal to flick their tail from the water was recorded using a stopwatch. The median result of three consecutive trials was recorded and reported for all animals tested.

### Open field recordings

Animals were recorded for thirty minutes in a novel environment made of 0.9 cm thick, white acrylic made in house (dimensions: 22.86 cm tall x 35.56 cm wide x 33.91 cm wide) using digital cameras placed above the open field (resolution 1920 x 1080 pixels, Spedal, ASIN B08BPFDYN7) and lit by bright white light (USB LED ring lights, 10 inch diameter, ASIN B07Q3471S2) from above at 16 and 196 hours following the final fentanyl dose. Experimenters observed the mice and recorded escape jumps. Video recordings were analyzed using EthoVision XT software (Noldus) to determine mobility, distance traveled, and time spent in the periphery of the open field compared to the center (defined by 50% of the floor area split between each).

### Tissue collection, RNA isolation, and immunohistochemistry (IHC)

For collection of RNA for sequencing, RiboTag-expressing mice were taken immediately after the last behavioral experiment, 16 hours following the last fentanyl or saline injection, and euthanized by decapitation using sharp scissors. The head was then immersed in ice cold phosphate buffered saline (PBS, 0.154 M sodium chloride, 0.141 M sodium phosphate monobasic, 0.1 M sodium hydroxide) for ~10-15 seconds and the brain then carefully extracted from the skull. Half of each brain was dissected following sagittal separation along the midline and the striatum was isolated and homogenized in supplemented homogenization buffer (50 mM Tris HCl, 100 mM MgCl2, 1% NP-40, 1 mM DTT, 1 x protein inhibitor cocktail [Millipore Sigma, catalog #P8340], 200 U/mL RNasin [Promega, catalog #N2611], 100 ug/mL cycloheximide [Millipore Sigma, catalog # C1988] and 1 mg/mL heparin, McKesson product # 1255341). All homogenates were centrifuged, and the supernatant was collected. From each sample, 10% of supernatant was set aside as the whole transcriptome sample (“input”, IN), and the remaining sample was processed to isolate ribosome bound RNAs (“immunoprecipitate”, IP see Fig S1A). All aspects of tissue processing, immunoprecipitation, and RNA-Seq library generation are described in detail in previous work from our group [44]. In the present study, RiboTag−/− animals were used to generate negative control samples. RNA sequencing and expression analysis cDNA libraries were prepared and sequenced as described in the Supplementary Methods. Validation of RNA expression was confirmed with quantitative PCR.

For collection of paraformaldehyde (PFA) fixed tissue, mice were injected with a fatal dose of Euthasol (pentobarbital sodium 390 mg/mL and phenytoin sodium 50 mg/mL, Virbac) diluted 1:1 with an equal volume of saline, according to their body mass (0.1 mL of diluted solution per 10 g). After reaching sufficient analgesic depth, the chest cavity was opened, the heart was pierced with a needle, and the body was perfused with ice-cold PBS to remove circulating blood and then by 4% PFA (40 g prills [Sigma product #441244] or powder [product #158127] per liter, prepared in PBS) to induce tissue fixation. The brain from each animal was carefully extracted from the skull following perfusion and post-fixed in 4% PFA for 16 hours at 4 °C and then transferred to a 30% sucrose solution (made in PBS as well). Sections were then sliced using a cryostat to 30 - 40 μm thick and stored in PBS with 0.01% sodium azide as a preservative (Millipore Sigma cat #S2002) until IHC was performed. Free floating sections were prepared for IHC by incubating in blocking buffer (4% normal donkey serum, 1% bovine serum albumin, 0.5% Triton-x 100, 0.5% Tween-20, 0.5% sodium deoxycholate) prepared in Tris-buffered saline (TBS, 0.137M sodium chloride, 0.01 M Tris base, pH to 7.4 with HCl) for 1 hour at room temperature using a nutating shaker. Sections were then moved to the primary antibody solution (blocking buffer with primary antibodies as described below) for incubation at 4 degrees C, shaking, for 72 hours. We used the following primary antibodies: Rabbit anti-Iba1 (1:500, Fujifilm Cellular Dynamics product #019-19741), Rat anti-C1q (1:1000, Abcam cat #ab11861), and Goat anti-PSD95 (1:500, Abcam catalog #ab12093). Following primary antibody incubation, sections were washed three times for five minutes each in blocking buffer followed by incubation in the secondary antibody solution for 1 hour, shaking at room temperature. We used the following secondary antibodies: Donkey anti-Rabbit AlexaFluor 555 (1:1000, Abcam cat #ab150062), Donkey anti-Rat AlexaFluor 488 (1:1000, Invitrogen catalog #A-21208), and Donkey anti-Goat AlexaFluor 647 (1:1000, Invitrogen catalog #A-21447). Following the secondary antibody incubation, sections were moved to TBS with one drop of DAPI solution (NucBlue Fixed Cell ReadyProbes Reagent, Invitrogen catalog #R37606) for thirty minutes, followed by five washes in TBS for ten minutes each before being mounted on slides using Prolong Diamond Antifade Mountant (Invitrogen catalog #P36965) or mounting media prepared in house (4.5 mL glycerol, 0.5 mL molecular grade biology water, 67 uL 1.5 M Tris pH 8, and 25 mg propyl gallate for 5.067 mL total).

### RNA sequencing and data analysis

We prepared cDNA libraries using the SMARTer Stranded Total RNA-Seq Kit v2 – Pico Input Mammalian (Takara, catalog # 634413). These were sent for quality control testing followed by sequencing using a NovaSeq two-lane chip on the Illumina system at the Northwest Genomics Center facilities at the University of Washington (https://nwgc.gs.washington.edu). The raw fastq files were concatenated (see supplemental materials for the Python script) to combine the data from two lanes, preserving separate forward and reverse runs from the paired-end samples. Quantification of transcripts was performed via Salmon software [45] using the mouse transcriptome downloaded from Ensembl (bash script also available in the supplemental materials). Determination of differentially expressed genes was performed using the R software DESeq2 package [21], followed by generation of gene modules based on Weighted Gene Coexpression Network Analysis (WGCNA) [23]. Briefly, the gene count matrix from the differential expression analysis for all IP samples that was generated using DESeq2 was filtered to remove zero-variance genes and genes below a minimum sequence count of five in at least two of the biological replicates. A signed topological overlap matrix was generated for clustering the genes. Module membership was assigned using a dynamic tree cut, and highly correlated modules were merged by reclustering module eigengenes. Gene module names were randomly assigned using the “Crayola” color palette to avoid introducing experimenter bias. Scripts for all RNA sequencing data processing and analysis are available in the supplementary materials.

### Quantitative PCR (qPCR) analyses

The mRNA expression levels for input (IN) and immunoprecipitated (IP) samples were assessed via qPCR on a Quant Studio Pro 7 using Power Sybr (ThermoFisher catalog #4389986). cDNA libraries were standardized across samples and targets with Ct values being normalized to that of peptidylprolyl isomerase A (PPIA). Primer sequences (see Table 1) were designed using the IDT PrimerQuest tool. Relative quantification and fold enrichment was obtained using the standard curve method [46].

**Table 1:**
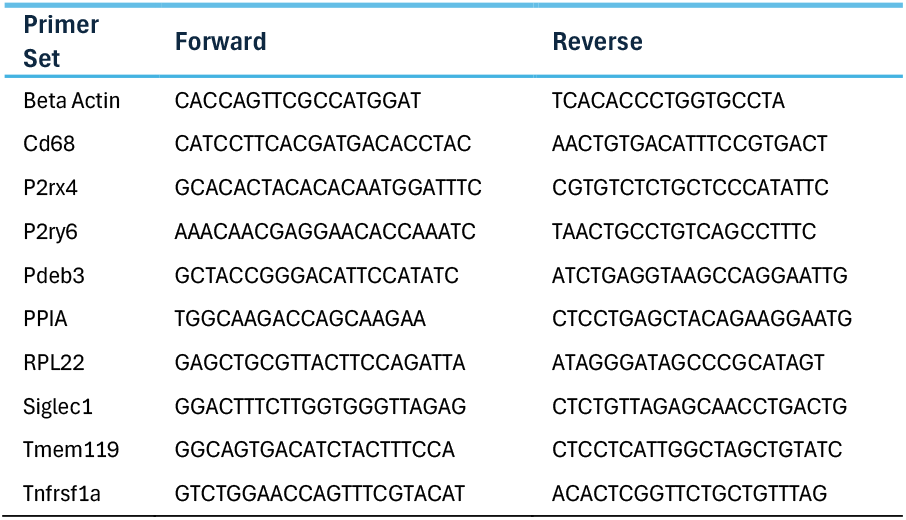
Primers Used for PCR.

**Table 2:** Mice represented in the RNA sequencing data.

| Drug | Withdrawal Cycles | N (sex = male, female) |
| --- | --- | --- |
| Saline | One | 10 (5, 5) |
| Fentanyl | One | 9 (4, 5) |
| Saline | Five | 4 (2, 2) |
| Fentanyl | Five | 7 (3, 4) |

### Confocal microscopy and image analysis

Following IHC, slices were imaged using either a Leica SP8 or a Nikon A1R confocal microscope. The SP8 imaging used an HC PL APO CS2 objective (20x, 0.75 NA). Slides were illuminated using a 470 – 670 nm tunable white light laser with power no greater than 30% for each channel. The average voxel size was 0.135 x 0.135 x 0.684 μm and an image size of 2048 x 2048 pixels (pixel dwell time 0.375 μs) using 4x line averaging. The A1R imaging used a CFI Plan Apo Lambda objective (40x, 0.95 NA). Slides were illuminated from a white light source with four filter cubes (DAPI, FITC, TRITC, and Cy5), with power no greater than 9 for each channel. Individual microglia reconstruction and quantification was performed using 3DMorph software [18].The average voxel size was 0.156 x 0.156 x 0.650 μm and an image size of 1024 x 1024 pixels using 16x line averaging. A bit-depth of 8 or 16 and a zoom factor of two was used for all images.

### Statistical methods and figure generation

We used Generalized Linear Mixed Models (GLMM) with JMP Pro 18. If there was no overall effect of sex, the groups were collapsed across sex, leaving treatment and cycle as fixed effects, and the mouse was treated as a random effect nested within treatment and cycle. Post-hoc comparisons were made using Tukey’s HSD test. RNA sequencing bioinformatics were performed in R, Python, and MATLAB.

## Code and Data Availability

Code for RNA-Seq pre-processing, differential expression analysis, qPCR analysis, and behavioral analysis is hosted at: https://github.com/rentazillla/OneVersusFive. Code for WGCNA and overrepresentation figure generation and analysis is hosted at: https://github.com/DrCoffey/Five-Cycle-WGCNA-Manuscript. Raw RNA-Seq data files are available upon reasonable request.

## Acknowledgments

Supported by R01DA052618, T32DA007278, R00DA052571.

Special thanks to Atom Lesiak for support with preparing the RNA for sequencing, to the UW Northwest Genomics Center for their cDNA library quality testing and for sequencing the RNA isolated in this study, to David Cook and Mayumi Yagi for use and training on the Leica SP8 confocal microscope, to Daryl Hackney for use and training on the Nikon A1R confocal microscope, and to Michele Kelly for support with mouse husbandry, statistical methods, and review of the manuscript. We thank the NIDA Drug Supply Program for supplying fentanyl.

## Author Contributions

David J. Bergkamp (performed experiments, analyzed data, wrote manuscript), Kevin R. Coffey (analyzed data, wrote manuscript), Aliyah J. Dawkins (performed experiments, analyzed data), Madelyn T. Rice (performed experiment, analyzed data), Ari M. Peden-Asarch (performed experiments, analyzed data), and John F. Neumaier (supervised all aspects of study, analyzed data, wrote manuscript).

---

**Figure S1.**
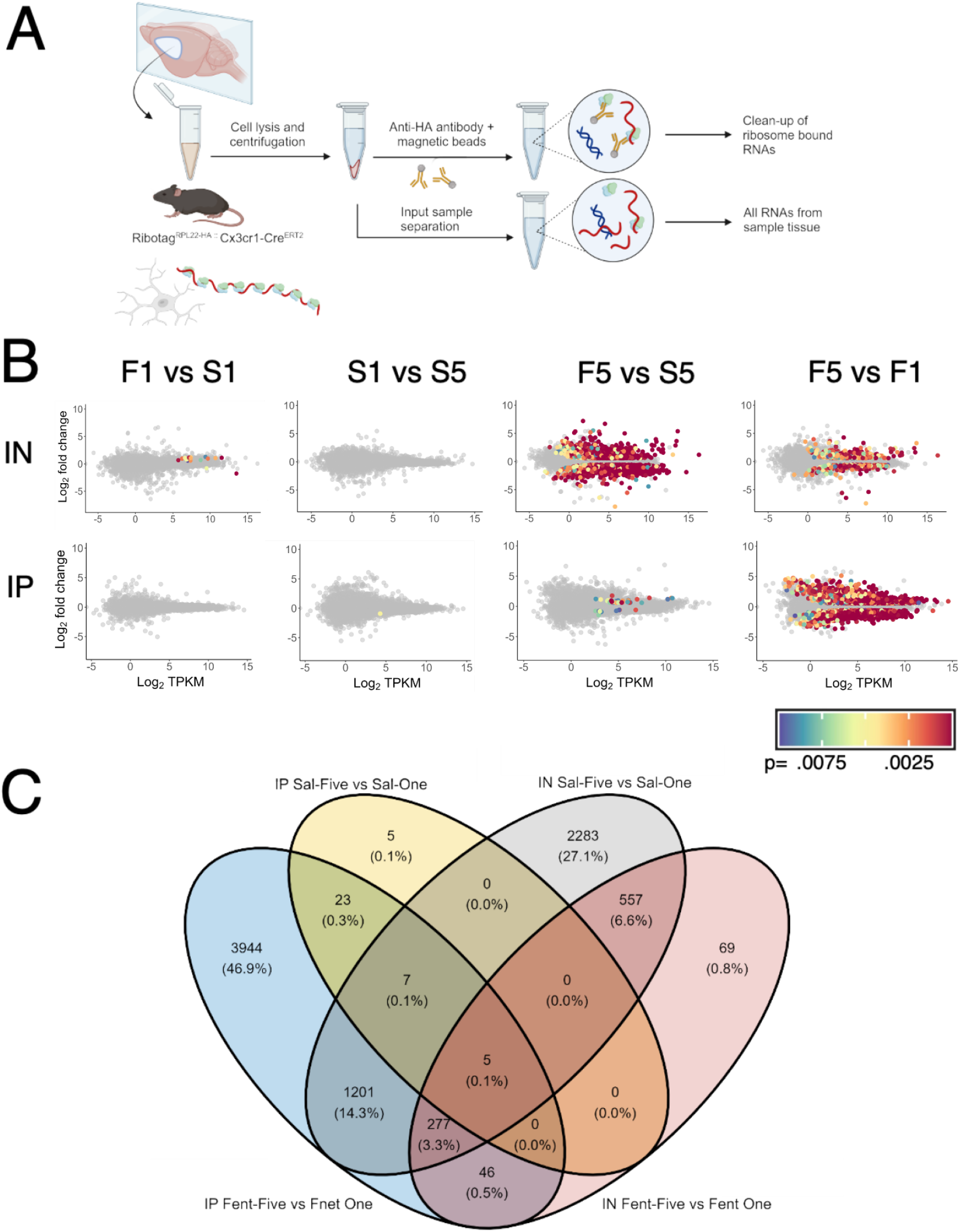
RNA Sequencing results. **A**, RiboTag RNA purification scheme. **B**, DESeq2 analyses show the comparisons within each Differentially expressed genes were defined as those with an adjusted p-value < 0.01 (method of Benjamini-Hochberg) and abs(Log2FoldChange) > 0.6. **C**, Venn diagram illustrates the number of DEGs in pair-wise comparisons between total striatal Input RNA (IN) and RiboTag IP RNA from Fent-Five vs. Fent-One (IP and IN, blue and red, respectively) and Sal-Five vs. Sal-One (IP and IN, yellow and grey, respectively). The largest number of DEGs was found in the IP Fent-Five vs. Fent-One comparison (blue ellipse, 5503 genes), and relatively few of these DEGs was also found in the other comparisons. The IN Fent-Five vs Fent-One RNA also contains the microglial translatome RNAs, so there is moderate overlap (blue and red ellipses, 328 genes). However, there are quite few DEGs shared between the IP samples from the Fentanyl DEGs and the Saline DEGs (blue and yellow ellipses, 35 genes).

**Figure S2.**
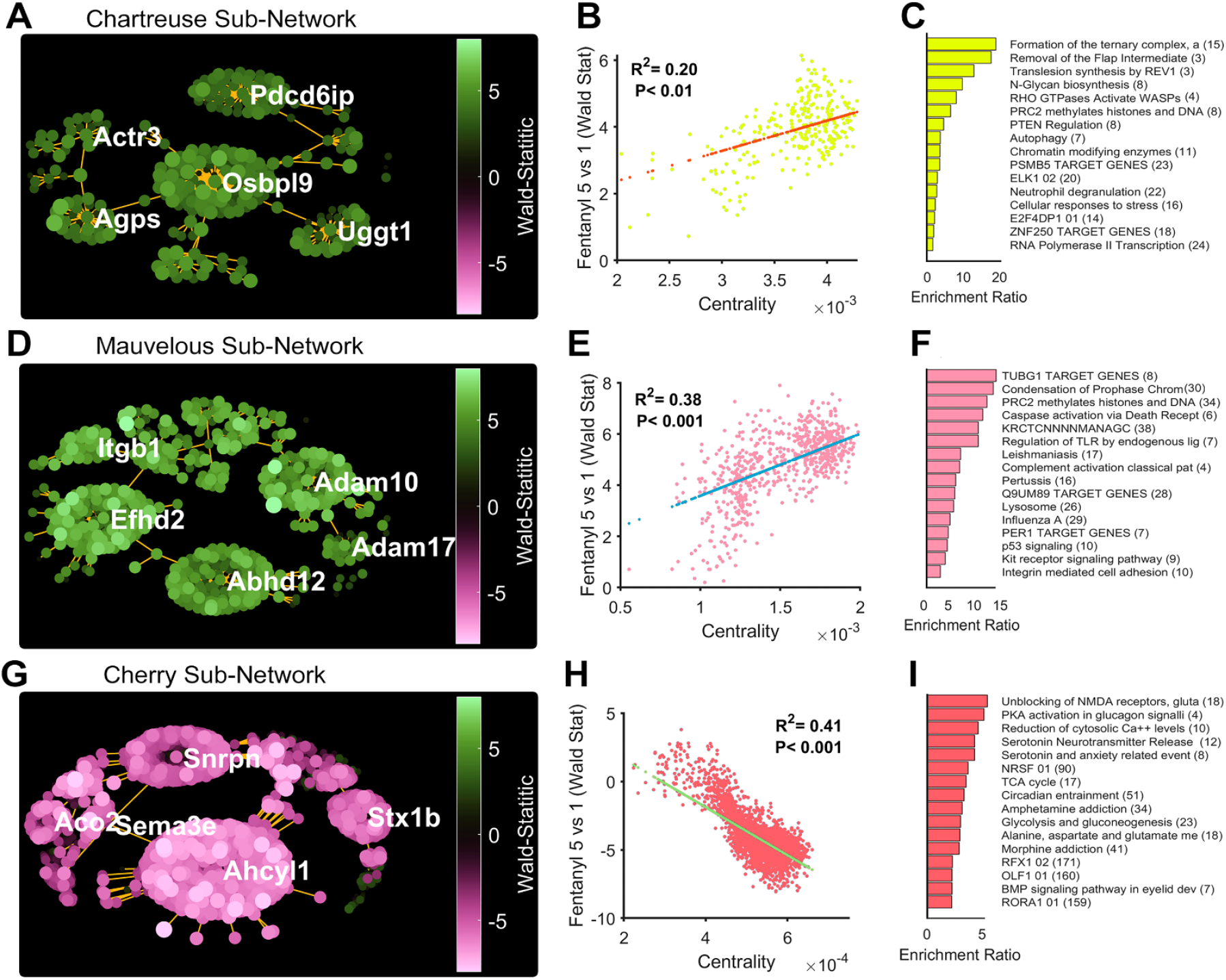
Additional gene modules with substantial overrepresentation of up or downregulated genes after repeated opioid withdrawal. Minimum spanning trees for genes in the Chartreuse (**A**), Mauvelous (**D**), and Cherry (**G**) modules; the Wald statistic for each point is indicated in the colored calibration bars. **B, E, H** show the correlation of each gene’s centrality to the corresponding module and the Wald statistic for the comparison of that gene’s differential expression using DESeq2, the Pearson correlation coefficient and p values are indicated. **C, F, I**, Overrepresentation analysis on the corresponding modules was performed in WebGESTALT as described in the Methods using several databases (Keeg, Reactome, Wiki Pathways, Transcription Factor Target). The top four upregulated and downregulated gene sets from each database are displayed with the number of genes from each module that overlapped with the gene set shown in parentheses. The enrichment ratio is the number of genes from the corresponding gene set present in the module divided by the number of genes expected by chance.

